# Natural variation in seed coat color in lettuce and wild *Lactuca* species

**DOI:** 10.1101/2024.06.27.600409

**Authors:** Sarah L. Mehrem, Guido Van den Ackerveken, Basten L. Snoek

## Abstract

Seed coat color is a well described trait in lettuce (*Lactuca sativa*), varying from black to pale white pigmentation. In this study, we delve into seed coat color variation of several species within the *Lactuca* genus, encompassing *L. sativa* and 15 wild varieties, offering broader insights into the diversity of this trait. To capture seed coat color quantitatively, we use grey pixel values from publicly available images, enabling us to measure seed coat color as a continuous trait across the genus. Darker seed coats predominate within the *Lactuca* genus, with *L. sativa* displaying a distinctive bimodal distribution of black and white seed coats. *Lactuca virosa* exhibits the darkest seed coat coloration and less variation, while *Lactuca saligna* and *Lactuca serriola* display lighter shades and greater variability. To identify the polymorphic loci underlying the observed variation we performed GWAS on seed coat color in both *L. sativa* and *L. serriola*. For *L. sativa*, we confirmed the one known major QTL linked to black and white seed coat color, which we reproduce in two independent, published genotype collections (n=129, n=138). Within the same locus, we identify additional candidate genes associated with seed coat color. For *L. serriola*, GWAS yielded several minor QTLs linked to seed coat color, harboring candidate genes predicted to be part of the anthocyanin pathway. These findings highlight the phenotypic diversity present within the broader *Lactuca* genus and provide insights into the genetic mechanisms governing seed coat coloration in both cultivated lettuce and its wild relatives.

## Introduction

Seed characteristics have been a key factor in crop domestication, including changes in seed size, yield, dispersal or color [1]. Seed coat color is an important agronomic trait for plant seeds that are either consumed or processed for consumption, like soybean (*Glycine max*), maize (*Zea mays*), squash (*Curcubita maxima*), pinto beans (*Phaseolus vulgaris*) and rapeseed (*Brassica napus*) [2-6]. Many studies have explored the biology and genetics of seed coat pigmentation and variation in these crops and other species, including *Arabidopsis thaliana* [2, 4, 6, 7]. In plants, flavonoids like proanthocyanins and anthocyanins produce the dark seed coat pigmentation by accumulating within the seed coat (testa) [8]. In *Arabidopsis thaliana*, seed coat pigmentation is controlled via transcription factor complexes, formed by *basic helix-loop-helix* (*bHLH*) proteins with both *myeloblastosis* (*MYB*) and *WD-repeat* (*WDR*) proteins that directly regulate biosynthetic and transporter genes of the flavonoid pathway [7, 9-11]. The genetic model of one or more transcription factors directly regulating seed coat color formation through the flavonoid pathways has been proposed in various crop species. Genes belonging to the *MYB* and *bHLH* protein families are some of the most likely candidates [3-6, 12-14].

Lettuce (*Lactuca sativa, L*.) is a globally important leafy vegetable crop [15]. Since its domestication from *Lactuca serriola* around 4000 BC, lettuce has diversified through cultivation [16, 17]. Today, numerous Lettuce cultivars exist, grouped into seven horticultural types. The domestication of lettuce has resulted in significant changes in many traits, including increased seed size, non-shattering seeds, lack of spines on leaves, and delayed bolting [2,3]. These altered traits made lettuce more fit for human consumption and contributed to its widespread cultivation around the world [3]. Wild lettuce species have been the major source for introducing novel agronomic plant traits. Three germplasm pools, consisting of 20 *Lactuca* species, have been identified as the main contributors of genetic variation during and after crop domestication [17]. Phylogenetic analyses of nuclear ribosomal DNA of *Lactuca* species revealed two distinct clades within the genus. The three germplasm pools, including *L. sativa*, are all included within the same clade [18].

In commercial lettuce, seed coat color has been used as a marker for marker assisted selection for disease resistance, climatic adaptation, and desirable commercial traits [19]. Furthermore, a darker seed coat color has been linked to higher germination quality [20]. Generally, seed coat color in *L. sativa* appears to be either dark brown /black, yellow or white. While this trait is very distinguishable and highly heritable [5] knowledge on the underlying genetics has been limited. Only recently, a locus responsible for white seed coat color in *L. sativa* was identified, revealing a *MYB* transcription factor (*LsTT2*) as the causal gene. This gene harbors an early termination codon, resulting in white seed color due to the lack of anthocyanins [21, 22].

In this study we describe seed coat color variation in *L. sativa* and 15 wild species from within and outside of its primary germplasm pools, exploring how this trait manifests within the genus. To capture detailed variations in seed coat color and facilitate cross-species comparisons, we use grey pixel values from seed images instead of categorical or ordinal scales. This method enhances resolution and precision in describing light to dark pigmented seed coats. Using GWAS we seek to identify further QTLs in *L. sativa* and *L. serriola* that may influence seed coat color variation. While *LsTT2* is the causal gene for a white seed coat variation, we hypothesize that additional candidate genes associated with the flavonoid pathway may contribute to the varying color intensities observed in dark seed coats across several *Lactuca* species.

## Methods

### Phenotyping

Seed pictures were obtained through the Centre for Genetic Resources Netherlands (CGN) [23]. Genotypes included were part of the Special_collection_CGNSC002. As a proxy for seed coat color (dark to light seeds), we measured grey values of seeds for each genotype using ImageJ version 1.53a [24]. Per genotype five or six independent seeds within the image were measured along the length and width axis. The mean grey pixel value was recorded separately for both width and length axes, resulting in two measured grey values for each of the five seeds per accession. These values were also used to estimate the heritability and variation in seed color. Further seed coat color phenotypes were collected from U.S. National Plant Germplasm System (https://npgsweb.ars-grin.gov/gringlobal/search) and CGN and matched via accession IDs as recorded by Zhang *et al*. 2017. These phenotypes were scored as a binary phenotype, “white” and “non-white”. For GWAS, values very averaged across all measurements per genotype.

### Heritability

Broad Sense Heritability (BSH) was estimated using base R (Version 4.2.2) and the script Heritability_estimate.R [25]. Seed coat color was fitted in a simple linear model using the lm function, with genotype as the predictor variable and phenotype as the outcome variable. ANOVA was performed using the anova function on the fitted model to estimate variance components. To estimate broad sense heritability (*H*^2^) the following formula was used:

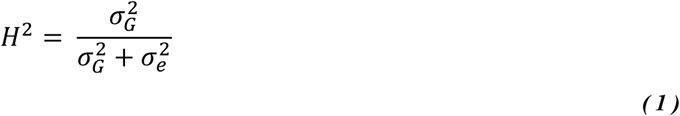

Where 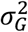 is the genotypic variance, 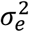 is the residual variance. 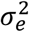 is estimated as the mean square of the residuals, as reported by ANOVA. 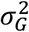 is estimated using the following formula:

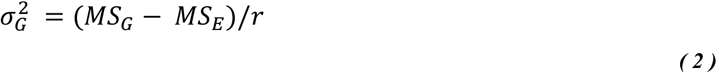

Where *MS*_*G*_ is the mean square of the predictor variable (Genotype), as estimated by ANOVA, *MS*_*E*_ is the mean square of the residuals and *r* is the number of replicates.

### Genotyping

For both the *L. sativa* (n=129) and *L. serriola* (n=199) population, single nucleotide polymorphism (SNP) data was obtained [17]. SNPs were filtered for a MAF > 0.05. a missing rate of <10% in *L. sativa* and *L. serriola* respectively for GWAS. Homozygous REF allele status was set as 1, heterozygous as 2 and homozygous ALT allele as 3. Missing alleles were set to a status of 2 (heterozygous). For the second SNP set we obtained imputed SNP data on a non-overlapping population of 214 *L. sativa* accessions, of which 138 accessions also had a recorded phenotype [26]. The imputed SNP set was filtered by removing duplicate SNPs, filtering for a MAF > 5% and a missing rate of < 25%. Homozygous wildtype status was set as 0, heterozygous as 0.5 and homozygous ALT allele as 1.

#### *Lactuca* sp. tree

The tree depicting *Lactuca* sp. taxonomy was curated manually from multiple phylogenies to place all *Lactuca* species that are phenotypically described in this study. For the basis, we used a coalescence-based phylogeny of single-loci nuclear genes of 12 *Lactuca* species [17]. *L. dregeana* was removed from the phylogeny because there was no phenotypic data available. The following species were added based on nuclear ribosomal DNA phylogenies [18]: *L. quercina, L. raddeana, L. biennis, L. floridana. L. canadensis* was placed into the tree, based on nuclear ribosomal DNA internal transcribed spacer phylogenies [27]. The *Lactuca* tree was then annotated based on reported germplasm pools that underly genetic variation of *L. sativa* [17]. *L. georgica* was changed to the secondary germplasm pool based on [28]. The tree was further annotated into two clades [18]. For visualizing the tree, the R package ggtree (Version 3.6.2) was used [29].

### GWAS

For Genome Wide Association Studies (GWAS) we used R version 4.2.2, the lme4QTL package and the scripts GWAS_lme4QTL.R and Zhang_etal_2017_seed_color_GWAS_2023.R [25, 30]. GWAS was performed on the filtered SNP set using a linear mixed model (relmatLmer and matlm), including the first five principle components (PCs) of the genotypes as a fixed effect and the kinship matrix as a random effect. The SNP as a covariate for a second GWAS was added in the matlm command as a fixed effect, PCs were fitted in the relmatLmer command. Kinship was calculated as the covariance matrix on the filtered SNPs. For multiple testing correction we used the Bonferroni threshold of 0.05. Linkage disequilibrium (LD) was calculated using the gl.report.ld command of the dartR R-package (Version 2.9.7), and the Pairwise_LD.R script [31]. To prune the LD matrix, the tagSNP function of the hscovar R-package (Version 0.4.2) was used, with a 0.9 pruning threshold, documented in the FigureS2_LDPlot.R script [32]. For additional GWAS in *L. serriola* we used the GAPIT3 package in R [33]. We ran GWAS with the multi-locus models “MLMM”, “FarmCPU”, “Blink” and single-locus model “SUPER”, as documented in script GapitGWAS_Serriola.R. Three PCs were included in the GWAS and a threshold for MAF > 0.5 was applied. Manhattan plots were visualized using the R packages ggplot2 (Version 3.4.4), cowplot (Version 1.1.1) and patchwork (Version 1.1.3) [34-36]. Quantile-Quantile (QQ) plots were created using the ggfastman R package [36]. Candidate genes were selected based on whether a significantly associated SNP was located within them, whether they were located in a region under high LD (R2 > 0.6) around the lead SNP, or whether the genes were predicted to be related to the flavonoid pathway and located in a 1Mbp window around that SNP. The genome annotation and gene function prediction of the Salinas reference genome V8 were used for annotation of candidate genes [37].

## Results

### Phenotypic distribution of seed color in *Lactuca* species

Seed coat color in *Lactuca* is diverse and shows species specific variation (**Figure 1A, Table S1**). The occurrence of black and white seeds is evenly distributed between *L. sativa* horticultural types, except for oilseed type varieties in our collection, which all have a dark seed coat (**Figure 1B**). To investigate the relation between seed color and evolutionary distance we combined a phylogenetic tree of the *Lactuca* species under study and their seed coat color, revealing a phenotype-specific relation with evolutionary distance (**Figure 1C**). Most *Lactuca* species have darker seed coat colors compared to the average *L. sativa* seed color. In general, species from germplasm pool I, which include *L. sativa, L. serriola, L. sagittata* and *L. altaica*, have lighter seeds coats compared the other germplasm pools and more distantly related *Lactuca* species. Comparing individual species, *L. virosa* has very dark, almost black, seeds, whereas *L. saligna, L. serriola* and *L. sativa* seeds appear dark to light brown. The largest within species variation in seed color was found in *L. sativa*, which strikingly showed a bimodal distribution, with more than half of the studied accessions having much lighter seeds than any other *Lactuca* species, including its closets relative *L. serriola*.

**Figure 1:**
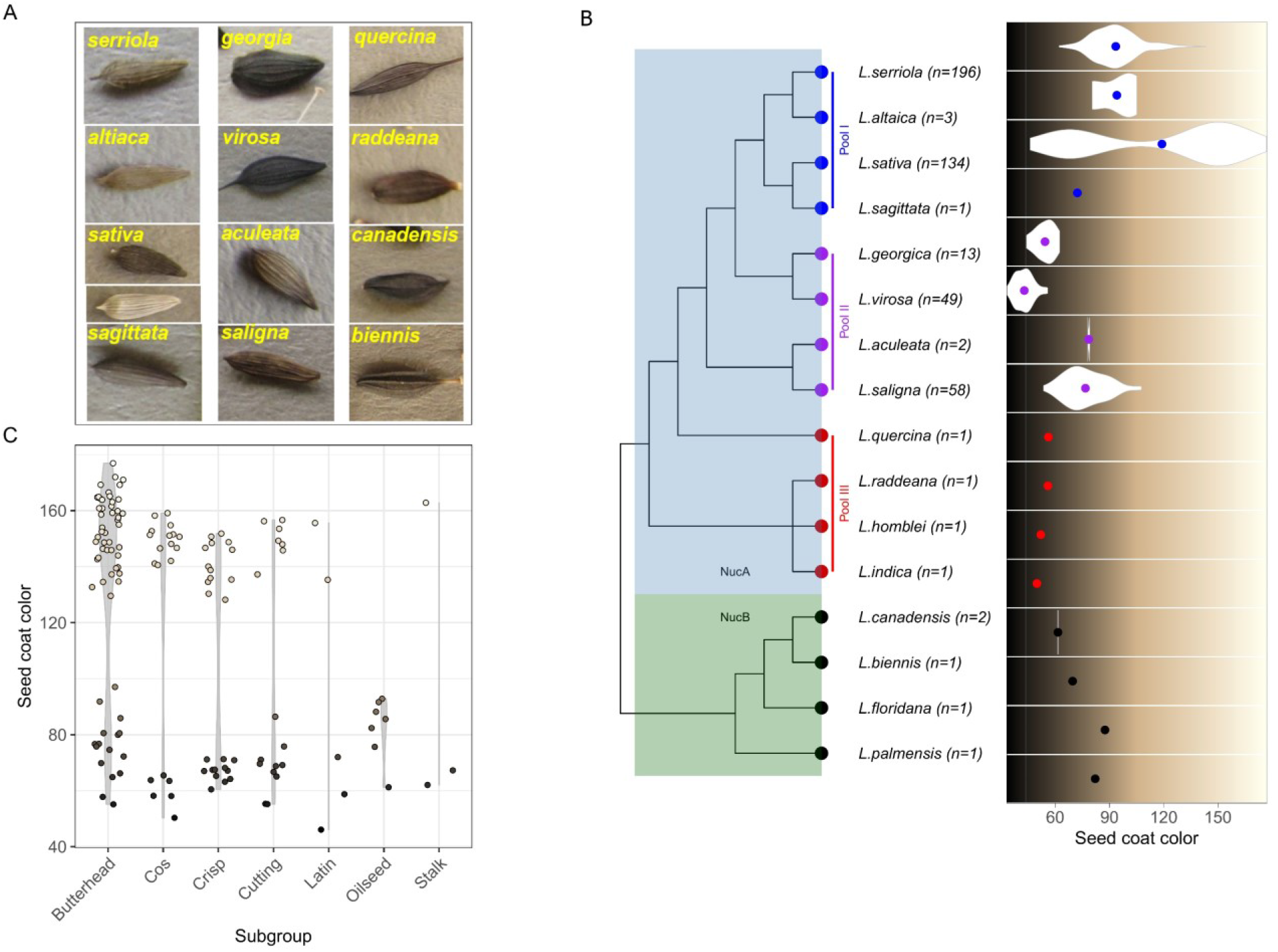
Distribution of seed coat color in *Lactuca* species. **A)** Seed pictures of *Lactuca* species. **B)** Seed coat color variation measured as grey value across the phylogenetic tree of *Lactuca* species. High grey values correspond to lighter seeds, low grey values to darker seeds. Species are indicated by their name. Nuclear genome pool (from Jones *et al*. 2018) is indicated by the blue and green background square. Germplasm pools according to Wei *et al*. (2021) are shown by the colored dots attached to the tree, blue: germplasm pool I, purple: germplasm pool II, red: germplasm pool III, black: outside the lettuce germplasm pool. Seed coat color variation is shown on the right in violin plots or single points. The points indicate the average seed coat color value (greyscale), violin plots show the seed coat color distribution per species. **C)** Seed coat color in *L. sativa* by horticultural type.

We estimated broad-sense heritability (BSH) for both *L. sativa* and *L. serriola* from raw seed coat measurements to assess the contribution of the genotype on the total phenotypic variability (**Table S2**). For *L. sativa* we found a BSH of 93% showing that most of the phenotypic variation can be explained by differences in genotype. For *L. serriola* the BSH was lower (40%), suggesting that genotypic variation contributes less to the observed variation in seed color.

### GWAS of the seed color trait in *L. sativa*

To identify genetic loci involved in seed color variation in *L. sativa* we performed GWAS (n =129). From the single SNP GWAS on the seed color scored as grey values, we found two quantitative trait loci (QTL). One QTL on chromosome 6 at 45,764,026 bp (-log_10_(p) > 18) and QTL on chromosome 7 at 50,317,920 bp (-log_10_(p) > 50; **Figure 2A, Table S3**). The locus on chromosome 6 cannot be decoupled from the locus on chromosome 7, as its significance diminishes when the top SNP on chromosome 7 is included as a cofactor in a second iteration of the GWAS (**Figure S1**). This indicates a possible false positive association due to linkage. Based on linkage, the QTL on chromosome 7 spans from approximately 49.2 Mbp to 52.3 Mbp, with 78 genes found in this locus (**Figure S2, Table S4**). The alleles of the lead SNP show a positive correlation with the phenotype (**Figure 2B**). The causative gene *LsTT2* for white seed color in lettuce (Lsat_1_v5_gn_7_35020.1) is located within the candidate locus. In addition, there are two previously reported seed coat color related genes that map to that region, Lsat_1_v5_gn_7_34841.1, a putative *NF-YA8* transcription factor, and Lsat_1_v5_gn_7_34600.1, a putative lycopene cyclase.

**Figure 2:**
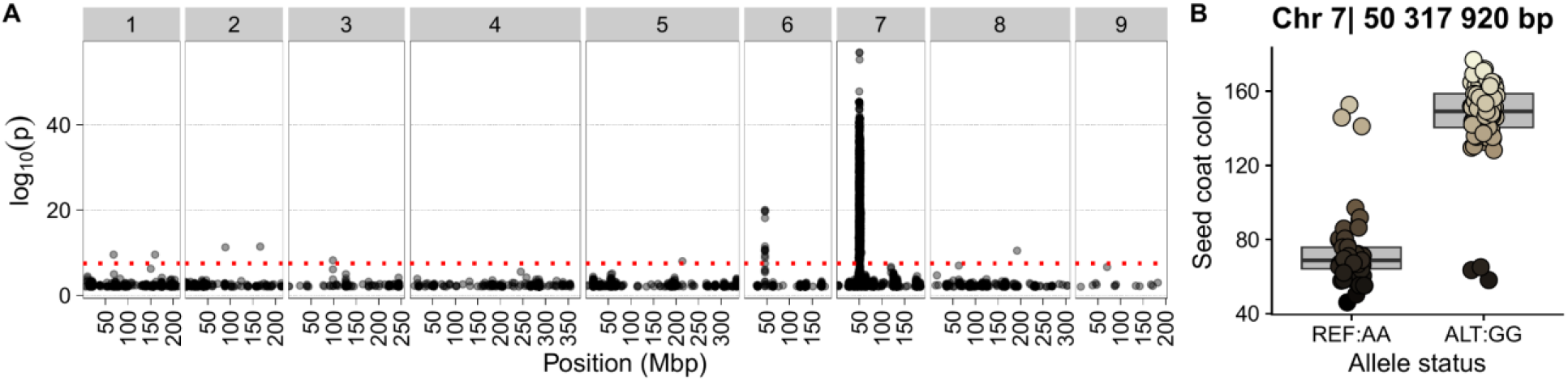
GWAS of seed coat color in *L. sativa*. **A)** Manhattan plot: Position in mega base pairs is shown on the x-axis, chromosome numbers are indicated on top, significance in – log10 (p) is show on the y-axis. Approximate Bonferroni threshold of -log10(p) > 7.4 is shown by the red horizontal line. **B)** Seed coat color distribution at the two alleles found for the most significantly linked SNP. Allele type, reference (REF) or alternative (ALT) show on the x-axis. Seed coat color proxy, grey value show on the y-axis. Color scale indicates the measured seed coat color

To determine if the variation in seed color in a different population would also be linked to the same QTL on chromosome 7 we performed GWAS on the *L. sativa* panel (n=138) from Zhang *et al*. 2017, of which only 5 accessions overlap with the *L. sativa* panel from Wei *et al*. 2021. The results indicate the same locus on chromosome 7 to be linked to the variation in seed color **(Figure S4**).

To identify QTLs linked to light or dark seed color variation respectively, we performed GWAS on both groups separately. This reduced the sample size considerably per GWAS (White seeds: 80 accessions, Black seeds: 49 accessions). No strong association for either phenotype is found (**Figure S3A+B, Table S5+6**). For white seed coat color, one SNP is significantly associated (-log10(P) > 7.8), located on chromosome 3 at 230,587,347bp (**Table S5**). Approximately 1.6 Mbp downstream of the SNP, a predicted *chalcone synthase 2* (*CHS2*) is located.

### GWAS of *L. serriola* seed coat color

To compare the genetic architecture of seed coat color of *L. sativa* with its wild relative *L. serriola* we performed GWAS on seed coat color in *L. serriola* (n=196). Using the same GWAS model applied to *L. sativa*, we identified two minor QTLs on chromosome 3 (QTL1) and 9 (QTL2), and one putative QTL on chromosome 5 (QTL3) (**Figure 3, Table 1, Table S7)**. For QTL1, we did not identify any candidate genes that could indicate a relation to seed coat color. Both SNPs of QTL2 fall within Lsat_1_v5_gn_9_101380.1, which is a putative calcium-dependent lipid-binding protein. Approximately 260kb downstream of QTL2, a predicted *gibberellin 3-beta-dioxygenase* (*GA3ox)* is the next potential candidate gene. We included a putative third locus (QTL3) as it showed a clear peak despite falling just under the significance threshold. The SNP is not located within a gene, but approximately 212 kb downstream of two copies of predicted *GA3ox*.

**Figure 3:**
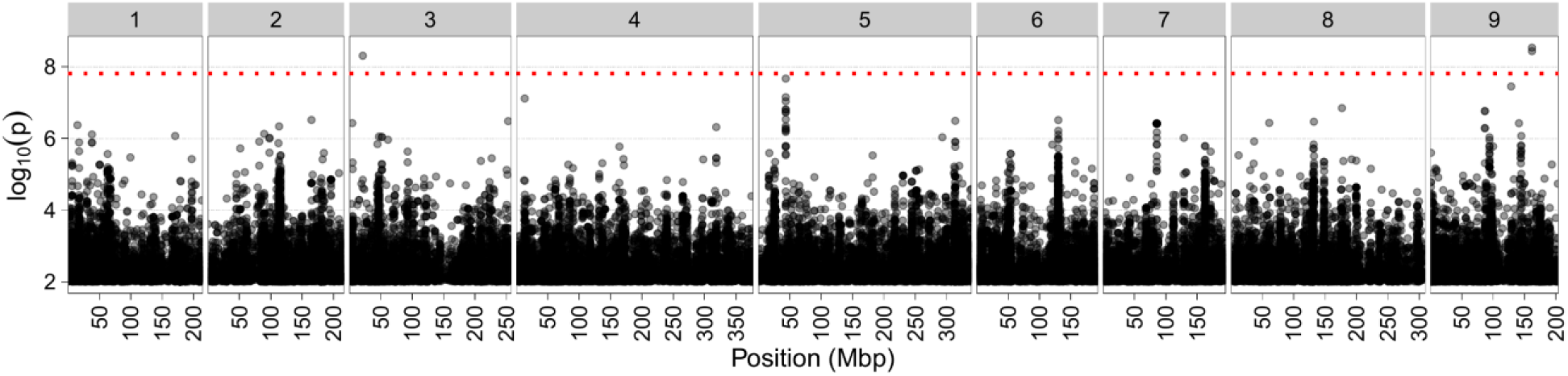
Manhattan plot of seed coat color GWAS in *L. serriola*. Position in mega basepairs is shown on the x-axis, chromosome numbers are indicated on top, significance in – log10 (p) is show on the y-axis. Approximate Bonferroni threshold of -log10(p) > 7.8 is shown by the red horizontal line.

**Table 1:** QTLs linked to seed coat color in *L. serriola*.

| Locus | Chr | Position (bp) | -LOG10(P) | Candidate genes | Functional annotation |
| --- | --- | --- | --- | --- | --- |
| QTL1 | 3 | 21,302,810 | 8,3 | - | - |
| QTL2 | 9 | 161,791,434 | 8,5 | Lsat_1_v5_gn_9_101380.1 | Calcium-dependent lipid-binding (CaLB domain) family protein |
|  |  |  |  | Lsat_1_v5_gn_9_101301.1 | hypothetical protein LOC100265167 \| K04124 gibberellin 3-beta-dioxygenase [EC:1.14.11.15] |
| QTL3 | 5 | 43,528,714 | 7,6 | Lsat_1_v5_gn_5_20961.1 | GA3ox1 \| gibberellin 3-oxidase (EC:1.14.11.15) \| K04124 gibberellin 3-beta-dioxygenase [EC:1.14.11.15] |
|  |  |  |  | Lsat_1_v5_gn_5_20940.1 | GA3ox1 \| gibberellin 3-oxidase (EC:1.14.11.15) \| K04124 gibberellin 3-beta-dioxygenase [EC:1.14.11.15] |

In the four additional GWAS conducted (BLINK, Super, MLMM, and FarmCPU), all but BLINK showed a consensus locus on chromosome 3, with one significantly associated SNP at 51,529,489 bp (**Figure S5, Figure S6, Table S8-11**). Approximately 150kb upstream lies one candidate gene (Lsat_1_v5_gn_3_40600.1), a putative anthocyanin 5-aromatic acyltransferase. None of the QTLs identified through the four additional GWAS models overlap with the results of lme4QTL. BLINK GWAS indicated a weak association that was not statistically significant (**Figure S5C**). For Super, the observed distribution of p-values suggests inflation (**Figure S6D**).

## Discussion

In this study we measured the seed coat color of a population of lettuce accessions and a panel of its wild relatives. We quantitatively described extensive variation in seed coat color both within and between species. Moreover, we identified one major QTL in *L. sativa* for seed color variation and several minor QTLs for seed color variation in *L* .*serriola*.

In *L. sativa*, the observed bimodal seed coat color distribution is a sign of human driven selection, and seed color itself functioned as a marker for breeding purposes, e.g. breeding for diseases resistance of climatic adaption [19]. Seed coat color has been linked to germination quality and disease resistance in the past, but as of now no conclusive results exist that validate this [20, 38]. The seed coat color trait in *L. serriola*, the ancestor species of *L. sativa*, shows an intermediate color, between the two modes, dark and white, of the bimodal distribution of *L. sativa*. This indicates that selection could have effectively shifted this trait towards the extreme phenotypes, leading to the clear bimodal distribution. Seed coat color has been used as a marker trait in recent history for *L. sativa*. However, its original significance during domestication and its relationship with traits such as seed oil content, given its initial cultivation as an oil crop, remain uncertain [17]. It is a common phenomenon in crops, that trait distribution changes through domestication, including non-target traits [39]. While not quantified in our dataset, *L. sativa* also shows yellow seed coat color, however it is reported to be rarer [19]. As grey values correspond to the darkness of the seed coat, variation in lighter colored seeds could be linked to yellow pigment, as those seeds also appear darker. While sample size for lighter seeds was relatively small, GWAS indicated one candidate locus for white seed coat color in *L. sativa*. As we scored the color quantitatively, we showed that there is pigment variation in lighter seeds. Near the candidate locus, a CHS2 gene is located, and CHS genes have been linked to lightness of white seeds [21].

We detected the already well described major QTL on chromosome 7 linked to seed coat color in *L. sativa* [21, 22, 40]. We validated this locus independently in a GWAS using SNPs from Zhang et al. (2017), which were called from different accessions with a minimal overlap of 5 accessions from Wei et al. (2021). We detect an additional locus on Chr 6; however, it is most likely a false positive association caused by linkage to the locus on Chr 7, and there are no genes found within this locus that could warrant further investigation. Phenotypes obtained for our study and the phenotypes obtained from the GRIN database were, to our knowledge, all conducted on independently grown plants. Given the striking similarity of the results, this further reinforces the previously reported very high heritability of the seed color trait.

Within the locus on chromosome 7, two additional genes could be further interesting candidates for non-white seed coat color variation. Predicted NF-YA8 has been found as an important candidate regulator of seed coat color intensity in rapeseed (*Brassica napus*) [41]. Predicted lycopene cyclase is a candidate for yellow seed coat color, as it is a key enzyme of the carotenoid biosynthesis pathway in *Arabidopsis thaliana*, and carotenoid concentration produces a yellow seed coat in soybean *(Glycine max)* [42, 43]. Compared to bimodal seed color variation in *L. sativa, L. serriola* shows a normal distribution of seed color variation and has a much lower broad sense heritability (BSH) than *L. sativa*. Likely the genetic architecture underlying this trait is more complex, or with more interaction with the environment, which also reduces the power for detecting QTLs for this phenotype using GWAS. Similarly, we do not detect any QTLs for dark seed coat color in *L. sativa*. In *L. virosa*, we observe the darkest seed coat, possibly due to high anthocyanin concentration, which presents a promising target for further investigation of dark seed coat color variation; GWAS or RNA sequencing could uncover whether this trait is linked to copy number variation that enhances anthocyanin biosynthesis.

Despite the reduced signal in *L. serriola*, we identified 3 minor QTLs. QTL 2 and 3 show promising candidates, as we identify 2 predicted Gibberellin-3-oxidases, which have been linked to darker seed coat color in barrel clover (*Medicago truncatula*) [44]. Using statistically more powerful GWAS models helped identify one additional candidate locus next to the single-locus GWAS in *L. serriola*, harboring a predicted anthocyanin 5-aromatic acyltransferase, which is part of the anthocyanin metabolic pathway in *Arabidopsis thaliana* [45].

Due to the domestication process, *L. sativa* has a reduced pool of genetic variation, compared to its progenitor *L. serriola* [17]. This can lead to strong association signals, with mono-genic or less complex genetics underlying the variation in many traits, such as seed coat color. While we have identified some intriguing candidate loci in *L. serriola*, the GWAS signals exhibit weak and indistinct peaks that do not overlap, raising concerns about potential false positives even with the stringent Bonferroni correction; further investigation with stronger evidence is needed. In addition, *L. serriola* SNPs were called against a *L. sativa* reference genome. While both species are genetically very similar to each other, they are not identical. If seed coat color in *L. serriola* is a more complex trait with multiple SNPs with small effect sizes, calling against a *L. serriola* reference genome would better represent the genetic variation within the species and potentially reveal more markers that would otherwise have been missed. We expect both species to also be different in terms of gene content, so even with a clear association in *L. serriola*, some of the causal genes might not be found on the *L. sativa* reference genome[26, 46].

While our study provides valuable insights, it is important to acknowledge several caveats. For some wild species, only one seed image was available at that time, making it difficult to fully compare seed coat color between species. While seed coat color correlates with grey values, reflecting anthocyanin content adequately, employing quantitative color identification methods would provide a more comprehensive phenotype description, accounting for the diverse pigments that contribute to color. Additionally, quantification of actual metabolites such as flavonoids or carotenoids would give a more accurate quantification of pigments in the seed coat, providing a suitable quantitative phenotype for GWAS. However, our study contributes by showcasing the efficacy of simple image analysis and using publicly available images to yield novel findings. Image resources for *Lactuca* are plentiful and can be scored for many phenotypes. Furthermore, by incorporating wild species, typically underrepresented, we want to emphasize the value of describing broader phenotypic diversity.

## Data availability

Scripts that were used for this study have been deposited at https://github.com/SnoekLab/Mehrem_etal_2024_SeedColor. Source data and supplementary tables are recorded in Supplement_SeedColor_V1.xlsx .

## Author contributions

BLS conceived the study, SLM, GvdA and BLS collected the data and phenotypes, SLM and BLS analyzed the data. SM and BLS wrote the manuscript with input from all co-authors.

## Acknowledgements

We thank the CGN for making the scores and images available. We thank Esther van den Bergh, Bram van Eijnatten and Sandra Smit for advice during writing.

## Supplement

**Figure S1:**
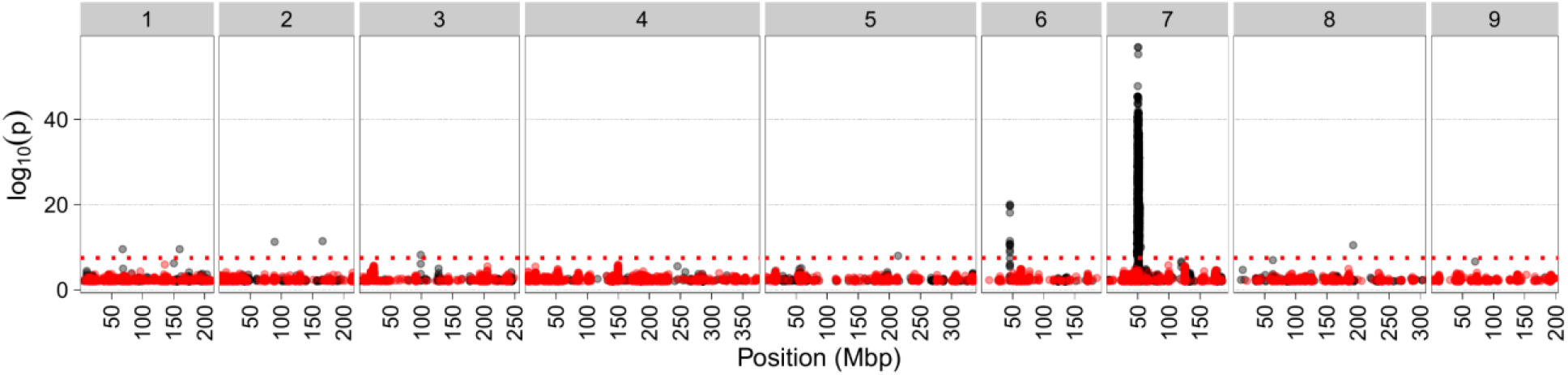
Manhattan plot of seed coat color GWAS in *L. sativa* with chromosome 7 lead SNP as fixed effect covariate. Position in mega basepairs is shown on the x-axis, chromosome numbers are indicated on top, significance in – log10 (p) is shown on the y-axis. Bonferroni threshold of -log10(p) > 7.4 is shown by the red horizontal line. Black dots are GWAS results without covariate, red dots are GWAS result with covariate.

**Figure S2:**
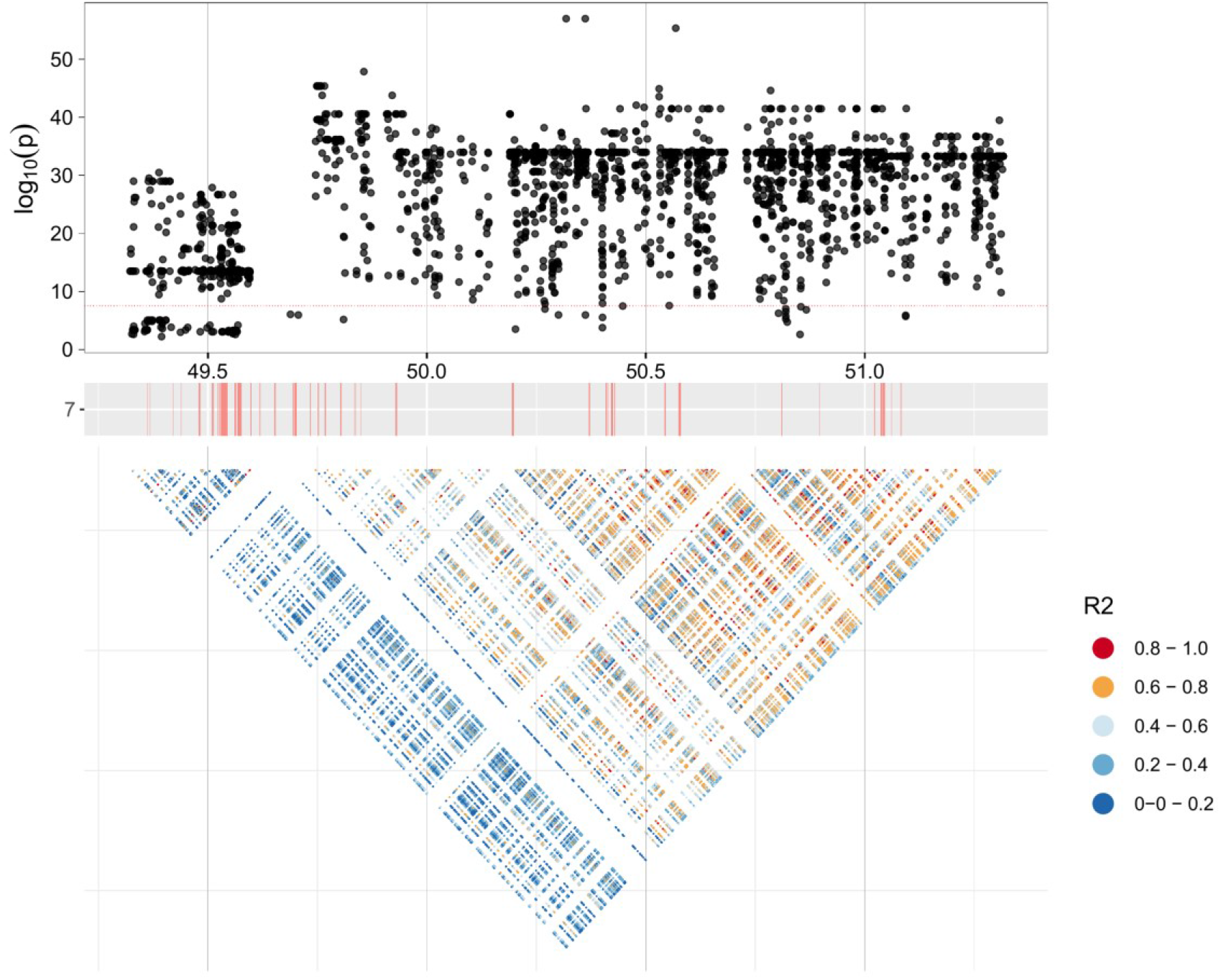
Seed coat color locus of *L. sativa* on Chr 7. Positions are shown in Mbp. Bonferroni corrected threshold is indicated by the red line. Significance is shown in -log10(P). Color corresponds to R2 value of LD calculation.

**Figure S3:**
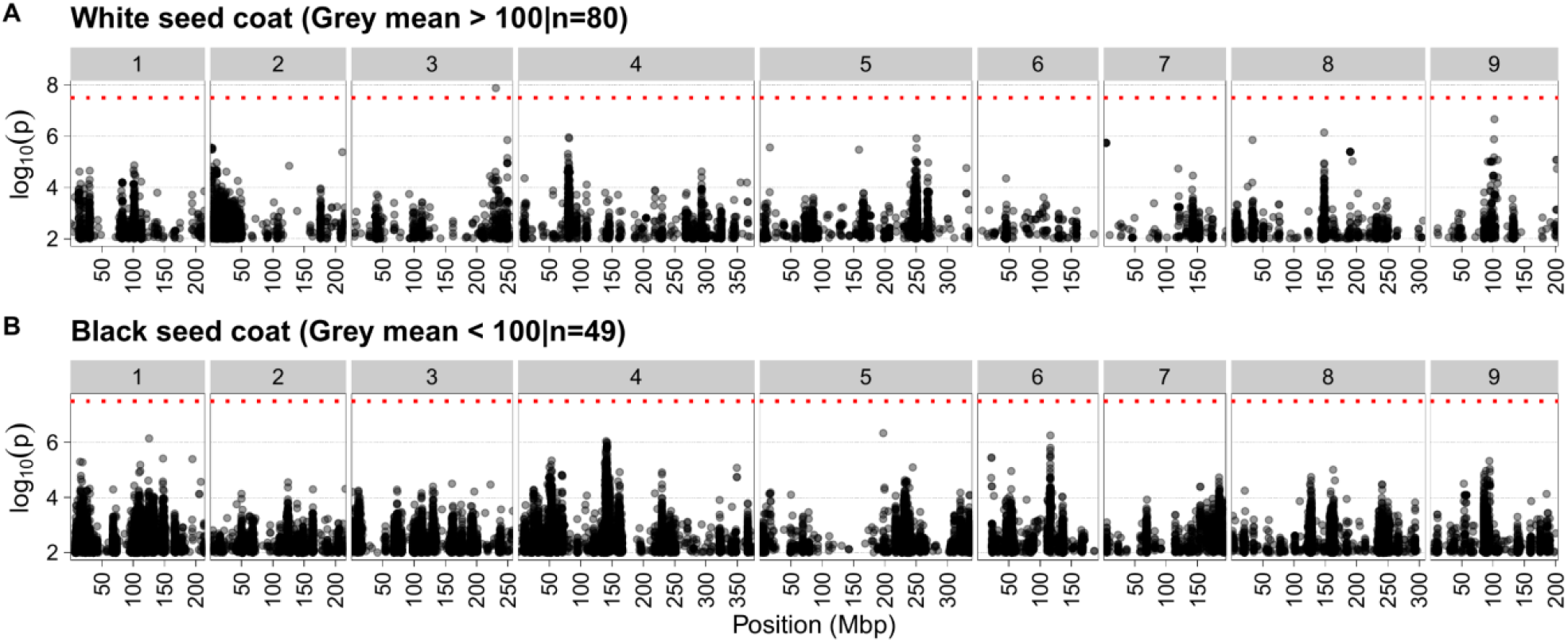
Manhattan plots of GWAS on black and white seeds. Position in mega basepairs is shown on the x-axis, chromosome numbers are indicated on top, significance in – log10 (p) is show on the y-axis. Bonferroni threshold of -log10(p) > 7.4 is shown by the red horizontal line. A) Manhattan of GWAS on white seeds. B) Manhattan of GWAS on black seeds.

**Figure S4:**
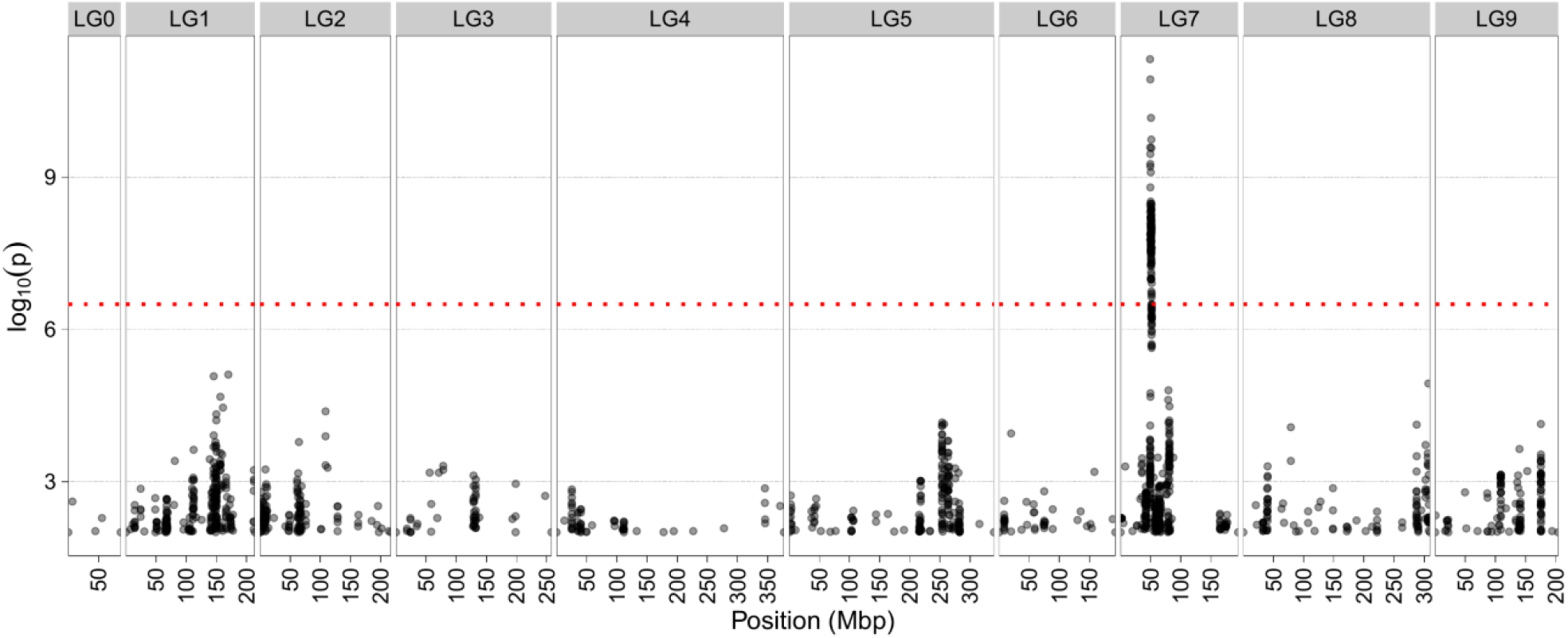
Manhattan plots of GWAS on seed coat color phenotype and SNPs from Zhang et al. (2017). Position in mega base pairs is shown on the x-axis, chromosome numbers are indicated on top, significance in – log10 (p) is show on the y-axis. Bonferroni threshold of -log10(p) > 6 is shown by the red horizontal line.

**Figure S5:**
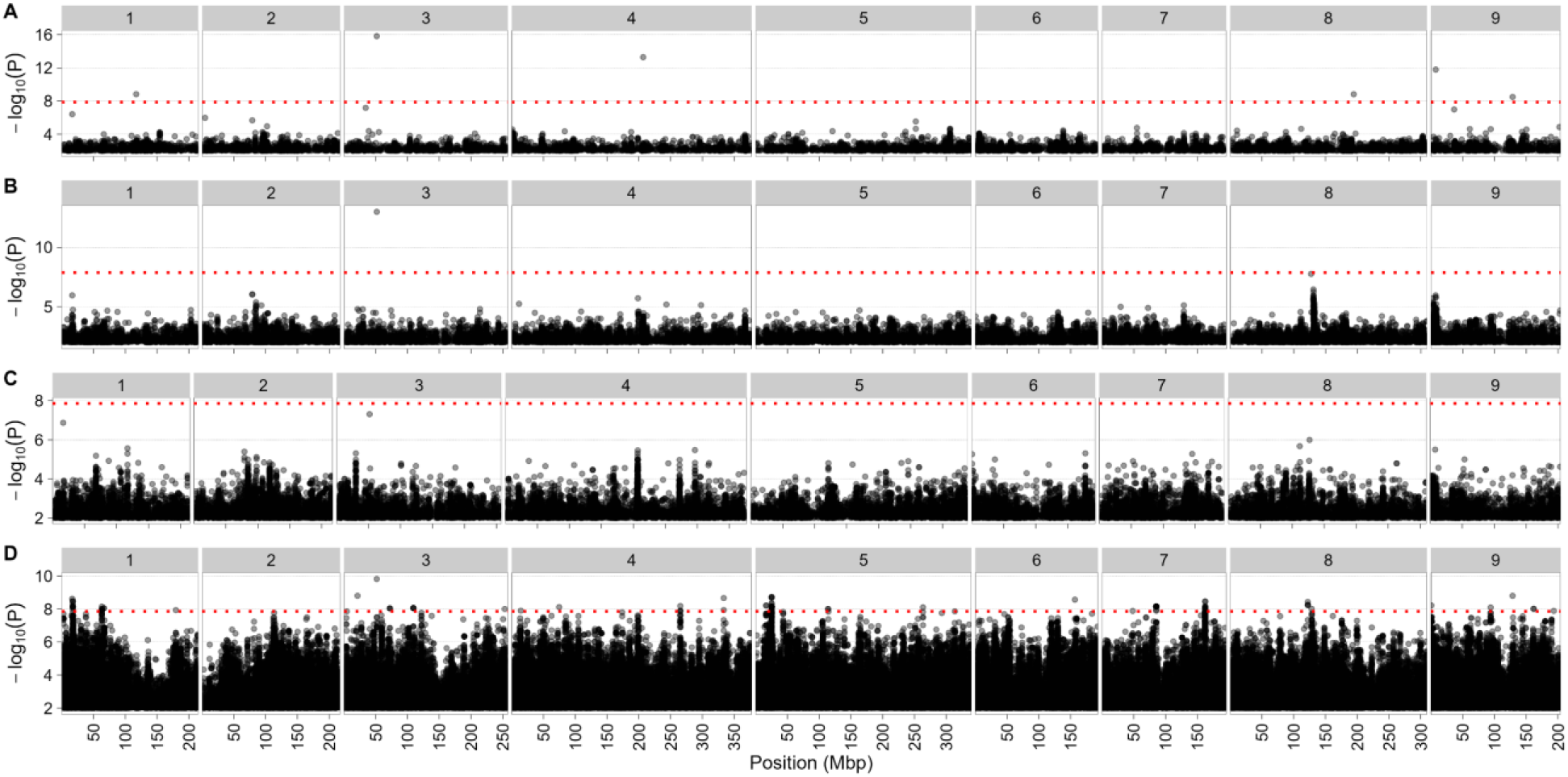
Manhattan plots of GWAS on seed coat color in *L. serriola*. A) FarmCPU, B) MLMM, C) BLINK, D) Super. Position in mega basepairs is shown on the x-axis, chromosome numbers are indicated on top, significance in – log10 (p) is show on the y-axis. Bonferroni threshold of -log10(p) > 7.8 is shown by the red horizontal line.

**Figure S6:**
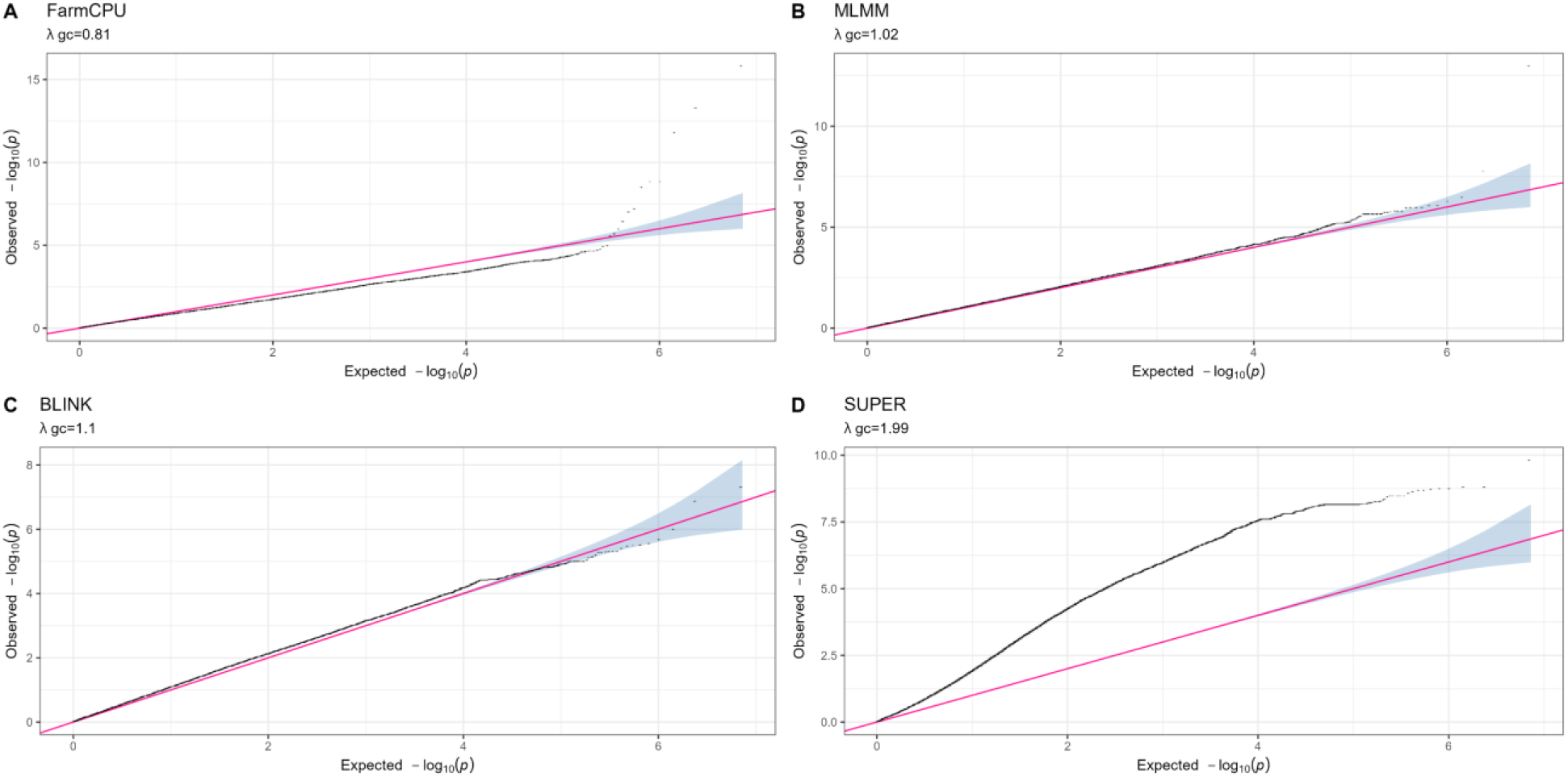
Quantile-Quantile (QQ) plots of GAPIT GWAS in *L. serriola* on seed coat color phenotype. A) QQ plot of FarmCPU. B) QQ plot of MLMM. C) QQ plot of BLINK. D) QQ plot of Super.

